# Using Transfer Learning for Automated Microbleed Segmentation

**DOI:** 10.1101/2022.05.02.490283

**Authors:** Mahsa Dadar, Maryna Zhernovaia, Sawsan Mahmoud, Richard Camicioli, Josefina Maranzano, Simon Duchesne

## Abstract

**Introduction:** Cerebral microbleeds are small perivascular haemorrhages that can occur in both grey and white matter brain regions. Microbleeds are a marker of cerebrovascular pathology, and are associated with an increased risk of cognitive decline and dementia. Microbleeds can be identified and manually segmented by expert radiologists and neurologists, usually from susceptibility-contrast MRI. The latter is hard to harmonize across scanners, while manual segmentation is laborious, time-consuming, and subject to inter- and intra-rater variabiltiy. Automated techniques so far have shown high accuracy at a neighborhood (“patch”) level at the expense of a high number of false positives voxel-wise lesions. We aimed to develop an automated, more precise microbleeds segmentation tool able to use standardizable MRI contrasts.

**Methods:** We first trained a ResNet50 network on another MRI segmentations task (cerberospinal fluid versus background segmentation) using T1-weighted, T2-weighted, and T2* MRI. We then used transfer learning to train the network for the detection of microbleeds with the same contrasts. As a final step, we employed a combination of morphological operators and rules at the local lesion level to remove false positives. Manual segmentations of microbleeds from 78 participants were used to train and validate the system. We assessed the impact of patch size, freezing weights of the initial layers, mini-batch size, learning rate, as well as data augmentation on the performance of the Microbleed ResNet50 network.

**Results:** The proposed method achieved a high performance, with a patch-level sensitivity, specificity, and accuracy of 99.57%, 99.16%, and 99.93%, respectively. At a per lesion level, sensitivity, precision, and Dice similarity index values were 89.1%, 20.1%, and 0.28 for cortical GM; 100%, 100%, and 1.0 for deep GM; and 91.1%, 44.3%, and 0.58 for WM, respectively.

**Discussion:** The proposed microbleed segmentation method is more suitable for the automated detection of microbleeds with high sensitivity.

## INTRODUCTION

Cerebral microbleeds are defined as small perivascular deposits filled from hemosiderin leaking from the vessels (Greenberg et al., 2009). They are recognized as a marker of cerebral small vessel disease, alongside white matter hyperintensities (WMHs) and lacunar infarcts (Wardlaw et al., 2013). Cerebral microbleeds are commonly present in patients with ischaemic stroke and dementia, and are more prevalent with increasing age (Roob et al., 1999; Sveinbjornsdottir et al., 2008; Vernooij et al., 2008). The presence of microbleeds has been linked to cognitive impairment and an increased risk of dementia (Greenberg et al., 2009; Werring et al., 2004).

*In vivo*, cerebral microbleeds can be detected as small, round, and well-demarcated hypointense areas on susceptibility-weighted (SWI) and T2* magnetic resonance images (MRIs) (Shams et al., 2015). Different studies have used various size cut-off points to classify microbleeds, with maximum diameters ranging from 5-10 mm, and, in some cases, a minimum diameter of 2 mm (Cordonnier et al., 2007). These are well correlated to histopathological findings of hemosiderin (Shoamanesh et al., 2011). In practice, microbleeds are labelled on MRI as being either “definite” or “possible” using visual rating (Gregoire et al., 2009). However, visual detection and segmentation is time consuming and subject to inter and intra rater variability, particularly for smaller microbleeds, frequently overlooked by less experienced raters. There is a need therefore for reliable and practical automated microbleed segmentation tools able to produce sensitive and specific segmentations at the lesion level.

Most of the microbleed segmentation methodologies currently available are semi-automated, i.e. they require expert intervention in varying degrees to produce the final segmentation (Barnes et al., 2011; Bian et al., 2013; Fazlollahi et al., 2015; Kuijf et al., 2013, 2012; Morrison et al., 2018). There have also been a few attempts at developing automated microbleed segmentation pipelines based on SWI scans, which in general have a higher sensitivity and resolution for microbleed detection (Dou et al., 2016; Hong et al., 2020; Roy et al., 2015; Shams et al., 2015; Van Den Heuvel et al., 2016; Wang et al., 2017; Zhang et al., 2018). These were shown to have sensitivity (ranging from 93 to 99 %) and specificity (ranging from 92 to 99%) at a neighborhood (“patch”) level, but less so at the voxel-wise, lesion level (Dou et al., 2016).

To address these issues, we employed techniques from the deep learning literature. In particular, convolutional neural networks (CNNs) have been successfully employed in many image segmentation tasks. Very deep convolutional networks such as ResNet (He et al., 2016), GoogLeNet (Szegedy et al., 2015), AlexNet (Krizhevsky et al., 2017), and VGGNet (Simonyan and Zisserman, 2014) have shown impressive performances in image recognitions tasks. ResNet50 has recently been used to detect microbleeds from SWI, achieving an accuracy of 97.46% at the patch level, outperforming other state-of-the-art methods (Hong et al., 2020).

As mentioned, most of these deep learning based studies only report patch-level results, without assessing their techniques on voxel-wise lesions on a full brain scan. The reported specificities are generally between 92-99% (Hong et al., 2020, 2019; Lu et al., 2017; Wang et al., 2017; Zhang et al., 2018). While high accuracy at a patch-level is important, when applied on the whole brain, even a specificity of 99% might translate into thousands of false positive voxels. In fact, applying different microbleed segmentation methods at a voxel level, Duo et al. report precision values ranging between 5-22%, in some cases leading to 280-2500 false positives on average (Dou et al., 2016). The proposed method by Duo et al. had a much better performance in terms of precision and false positives, with an precision rate of 44.31% and an average false positive rate of 2.74, however, their reported sensitivity was relatively lower (93.16%).

Thus, improving performance at the lesion level would be desirable. Further, given that SWI are not always collected in either clinical or research settings and/or are hard to harmonize in multi-centric settings, it would be useful if a more versatile algorithm was proposed, able to segment microbleeds from other, more general MRI contrasts (e.g. T1-weighted, T2-weighted, or T2*). To our knowledge, there is no published automated microbleed segmentation tool based on T1w/T2w/T2* acquisitions, which is the first contribution of our article.

The main challenge in developing automated microbleed segmentation tools using machine learning and in particular deep learning methods pertains to a general lack of reliable, manually labelled data. Our second contribution is how we address this problem by using tranfer learning to deal with the relatively small number of manually labelled microbleeds in our dataset. In light of promising results from other authors, we employed a pre-trained ResNet50 network, further tailoring it for another, relevant MRI segmentation task, namely the classification of cererbospinal fluid versus brain tissue, for which we were able to generate a large number of training samples. The pre-trained weights were then used as the initial weights for our microbleed segmentation network, allowing for a faster convergence with a smaller training sample.

A third contribution is how we employed post-processing to winnow out false positives. This strategy is described below, along with results on the performance of our technique, both at the patch and the pixel level. Transfer learning is a powerful paradigm that makes our algorithm potentially versatile on a number of MRI contrasts for microbleeds detection.

## MATERIALS AND METHODS

### Participants

Data included 78 subjects (32 women, mean age= 77.16 ± 6.06 years) selected from the COMPASS-ND cohort (Chertkow et al., 2019) of the Canadian Consortium on Neurodegeneration in Aging (CCNA; www.ccna-ccnv.ca). The CCNA is a Canadian research hub for examining neurodegenerative diseases that affect cognition in aging. Clinical diagnosis was determined by participating clinicians based on longitudinal clinical, screening, and MRI findings (i.e. diagnosis reappraisal was performed using information from recruitment assessment, screening visit, clinical visit with physician input, and MRI). For details on clinical group ascertainment, see Pieruccini‐Faria et al. 2021 (Dadar et al., 2021b, 2021a; Pieruccini‐Faria et al., 2021).

All COMPASS-ND images were read by a board-certified radiologist. Out of the whole cohort, participants in this study were selected based on the presence of WMHs on FLuid Attenuated Inversion Recovery MRIs as another indicator of cerebrovascular pathology and microbleeds on T2* images. Consequently, the sample was comprised of six individuals with subjective cognitive impairment (SCI), 30 individuals with mild cognitive impairment (MCI), six patients with Alzheimer’s dementia (AD), eight patients with frontotemporal dementia (FTD), seven patients with Parkinson’s disease (PD), three patients with Lewy body disease (LBD), five patients with vascular MCI (V-MCI), and 13 patients with mixed dementias. Given that ours is a study on segmentation performance, we assumed that there was no difference in the T2* appearance of a microbleed related to participants’ diagnosis.

Ethical agreements were obtained for all sites. Participants gave written informed consent before enrollment in the study.

### MRI Acquisition

MRI data for all subjects in the CCNA was acquired with the harmonized Canadian Dementia Imaging Protocol (www.cdip-pcid.ca; (Duchesne et al., 2019)). Table 1 summarizes the scanner information and acquistion parameters for the subjects included in this study.

**Table 1.** Acquisition parameters of the harmonized Canadian Dementia Imaging protocol.

| Sequence | Scanner Model | Matrix | Resolution (mm<sup>3</sup>) | Number of Slices | TR (msec) | TE (msec) | TI (msec) | Flip Angle |
| --- | --- | --- | --- | --- | --- | --- | --- | --- |
| T1w | GE | 256 × 256 | 1.0 × 1.0 × 1.0 | 180 | 6.7 | 2.9 | 400 | 11 |
|  | Philips | 256 × 248 | 1.0 × 1.0 × 1.0 | 180 | 7.3 | 3.3 | 935 | 9 |
|  | Siemens | 256 × 256 | 1.0 × 1.0 × 1.0 | 192 | 2300 | 2.98 | - | 9 |
| T2w | GE | 256 × 256 | 0.94 × 0.94 × 3.0 | 48 | 3000 | 11 | - | 125 |
|  | Philips | 256 × 254 | 0.94 × 0.94 × 3.0 | 48 | 3000 | 13 | - | 90 |
|  | Siemens | 256 × 256 | 0.94 × 0.94 × 3.0 | 48 | 3000 | 10 | - | 165 |
| T2* | GE | 256 × 256 | 0.94 × 0.94 × 3.0 | 48 | 650 | 20 | - | 20 |
|  | Philips | 256 × 256 | 0.94 × 0.94 × 3.0 | 48 | 650 | 20 | - | 20 |
|  | Siemens | 256 × 256 | 0.94 × 0.94 × 3.0 | 48 | 650 | 20 | - | 20 |

TR: repetition time; TE: echo time; TI: inversion time.

### MRI Preprocessing

All T1-weighted, T2-weighted and T2* images were preprocessed as follows: intensity non-uniformity correction (Sled et al., 1998), and linear intensity standardization to a range of [0-100]. Using a 6-parameter rigid registration, the three sequences were linearly coregistered (Dadar et al., 2018). The T1-weighted images were also linearly (Dadar et al., 2018) and nonlinearly (Avants et al., 2008) registered to the MNI-ICBM152-2009c average template (Manera et al., 2020). Brain extraction was performed on the T2* images using BEaST brain segmentation tool (Eskildsen et al., 2012).

### CSF Segmentation

The Brain tISue segmentatiON (BISON) tissue classification tool was used to segment CSF based on the T1-weighted images (Dadar and Collins, 2020). BISON is an open source pipeline based on a random forests classifier that has been trained using a set of intensity and location features from a multi-center manually labelled dataset of 72 individuals aged from 5-96 years (data unrelated to this study) (Dadar and Collins, 2020). BISON has been validated and used in longitudinal and multi scanner multi-center studies (Dadar et al., 2020; Dadar and Duchesne, 2020; Maranzano et al., 2020).

### Grey and White Matter Segmentation

All T1-weighted images were also processed using *FreeSurfer* version 6.0.0 (*recon-all-all*). *FreeSurfer* is an open source software (https://surfer.nmr.mgh.harvard.edu/) that provides a full processing stream for structural T1-weighted data (Fischl, 2012). The final segmentation output (aseg.mgz) was then used to obtain individual masks for cortical GM, deep GM, cerebellar GM, WM, and cerebellar WM based on the FreeSurfer look up table available at https://surfer.nmr.mgh.harvard.edu/fswiki/FsTutorial/AnatomicalROI/FreeSurferColorLUT. Since *FreeSurfer* tends to segment some WMHs as GM (Dadar et al., 2021c), we also segmented the WMHs using a previously validated automated method (Dadar et al., 2017b, 2017a) and used them to correct the tissue masks (i.e. WMH voxels that were segmented as cortical GM or deep GM by *FreeSurfer* were relabelled as WM, and WMH voxels that were segmented as cerebellar GM by *FreeSurfer* were relabelled as cerebellar WM).

### Manual Segmentation

The microbleeds were segmented by an expert rater (JM > than 15 years of experience reading research brain MRI) using the interactive software package Display, part of the minc-toolkit (https://github.com/BIC-MNI) developed at the McConnell Brain Imaging Center of the Montreal Neurological Institute. The software allows visualization of co-registered MRI sequences (T1w, T2w and T2*) in three planes simultaneously, cycling between sequences to acurately assess the signal intensity and anatomical location of an area of interest. Identification criteria was in accordance with Cordonnier et al., 2007, comprised a round area of hypointensity on T2* within the brain tissue and exclusion of colocalisation with blood vessels based on T1w and T2w information (Cordonnier et al., 2007). A maximum diameter cut-off point of 10 mm was used to exclude large hemorrhages (Cordonnier et al., 2007). No minimum microbleed size cut-off was used. Eight cases with varying number of microbleeds were segmented a second time by the same rater (JM) to assess intra-rater variability.

### Quality Control

We visually assessed the quality of the preprocessed images, as well as the BISON and FreeSurfer automated segmentations.

### Generating Training Data

*Transfer Learning CSF Segmentation Task:* 400,000 randomly sampled two-dimensional (2D) image patches were generated from the in-plane preprocessed and co-registered T2*, T2-weighted, and T1-weighted image slices. Half of the generated patches contained a voxel segmented as CSF by BISON in the center voxel of the patch, and the other half contained either GM or WM in the center of the patch. The patches were randomly assigned to training, validation and test sets (50%, 25%, and 25% respectively). To avoid leakage, patches that were generated from one participant were only included in the same set; i.e. the random split was performed at participant level (Mateos-Pérez et al., 2018). Figure 1 shows examples of the generated CSF and background patches.

**Figure 1.**
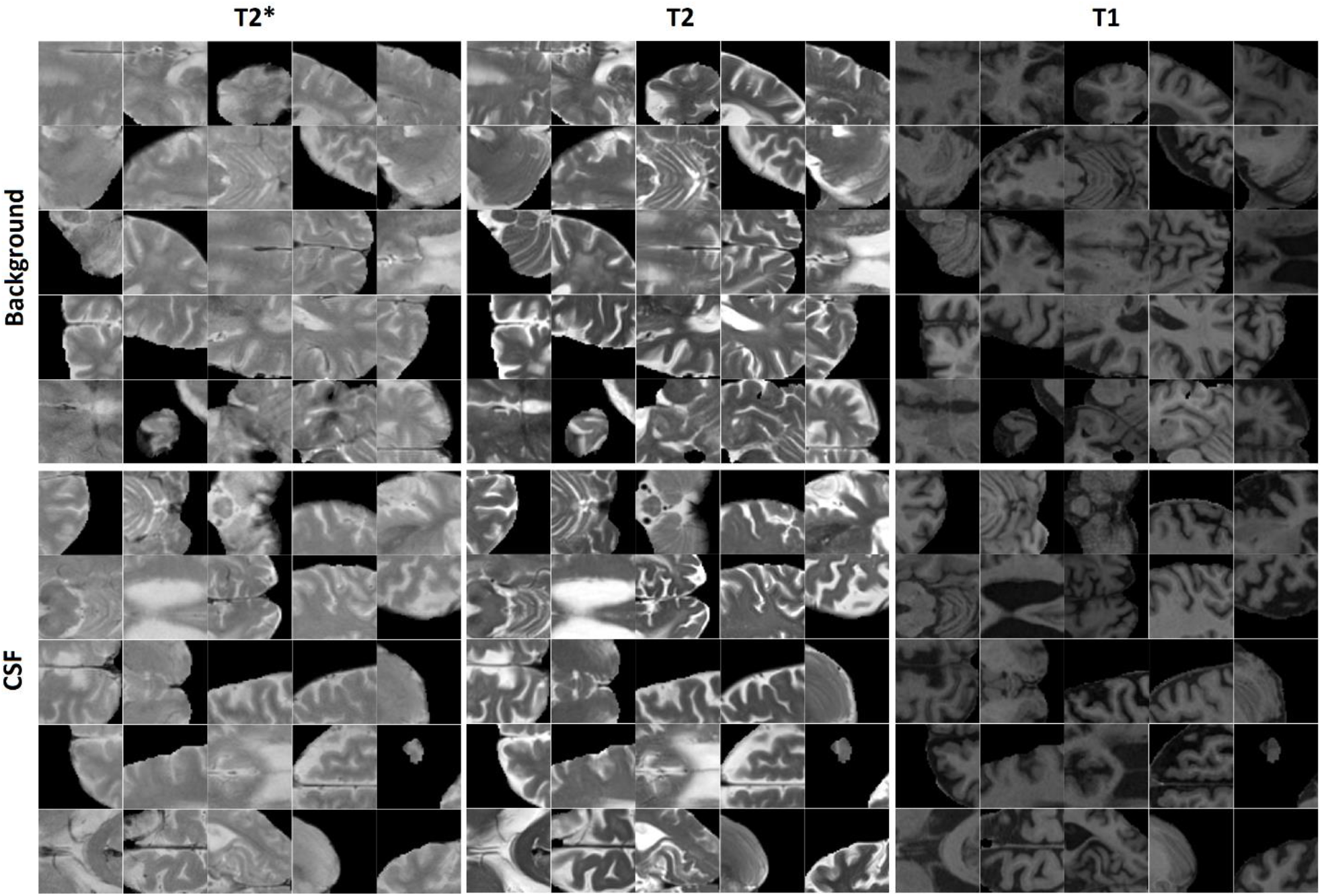
Examples of CSF and background patches generated for the CSF segmentation task.

*Microbleed Segmentation Task:* Similarly, 11,570 2D patches were generated from the preprocessed and co-registered T2*, T2-weighted, and T1-weighted in-plane image slices for microbleed segmentation. Half of the generated patches contained a voxel segmented as microbleed by the expert rater in the center voxel of the patch, and the other half was randomly sampled to contain a non-microbleed voxel in the center. The patches were randomly assigned to training, validation and test sets (60%, 20%, and 20% respectively) also at the participant level. We further ensured to include similar proportions of participants with small (1-4 voxels), medium (5-15 voxels), and large (more than 15 voxels) microbleeds in the three sets. Figure 2 shows examples of the generated microbleed and background patches.

**Figure 2.**
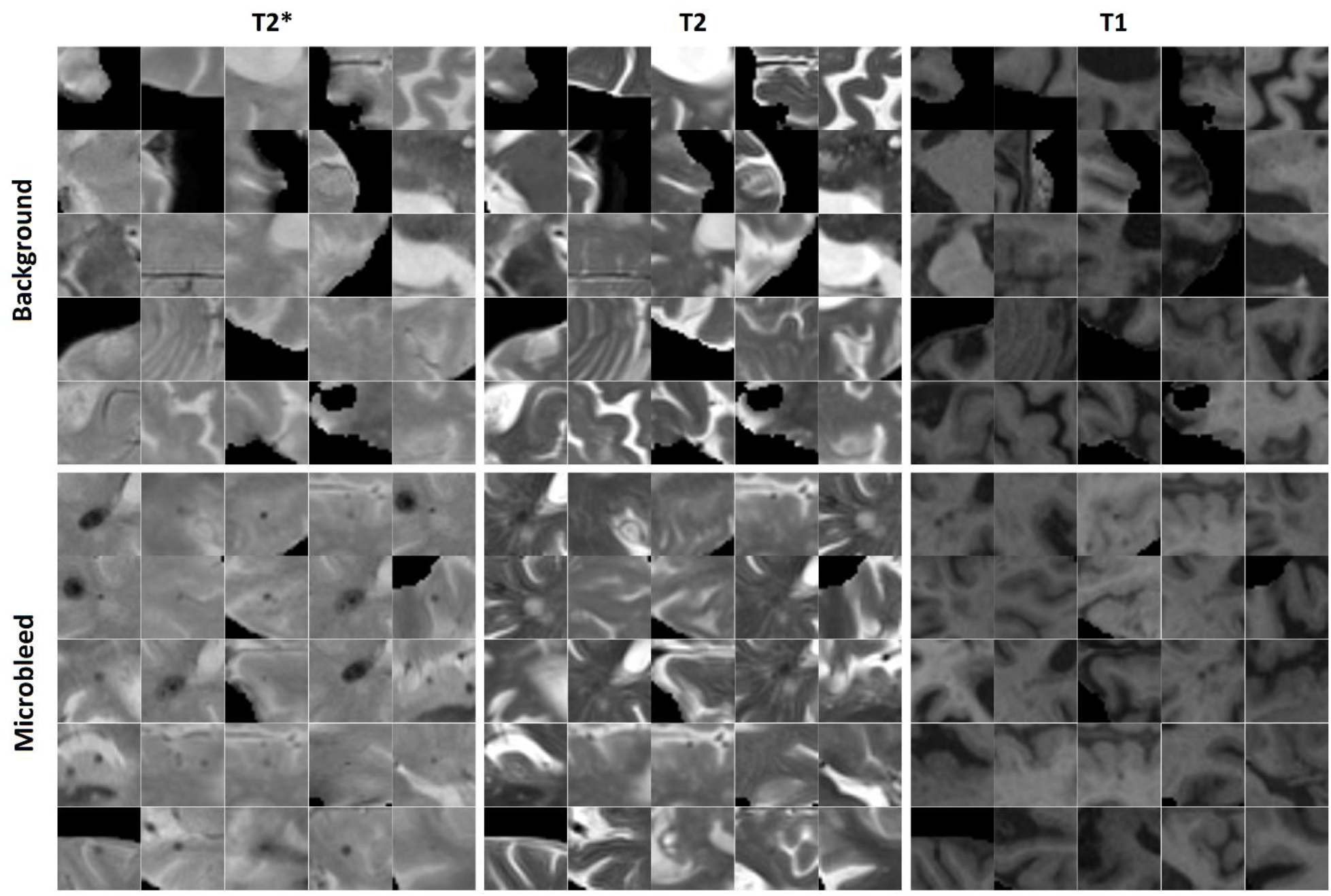
Examples of microbleed and background patches generated for the microbleed segmentation task.

We further augmented the microbleed patch dataset by randomly rotating the patches to generate additional training data. The random rotations were performed on the full slice (not the patches) centering around the microbleed voxel; therefore, the corner voxels in the patches include information from different areas not present in other patches. Matching numbers of novel background patches were also added to balance the training dataset. The performance of the model was assessed using the training dataset with no augmentation, and with adding 4, 9, 14, 19, 24, and 29 random rotations to the training set, respectively.

### Training the ResNet50 Network using Transfer Learning

We used the ResNet50 network (He et al., 2016), a CNN pre-trained on over 1 million images from the ImageNet dataset (Challenge, 2012) to classify images into 1000 object categories. This pre-training has allowed the network to learn rich feature representations for a wide range of images, which can also be useful in our task of interest. Our approach was to further train ResNet50 first on a task similar to microbleed segmentation (i.e. CSF vs. tissue) then on microbleeds identification itself.

To satisfy the input size requirements of ResNet50 network, all patches were resized to 224×224 pixels, and the T2*, T2-weighted, and T1-weighted patches were copied in to three channels to generate an RGB image (Figure 3). The last fully connected layer of ResNet50, which contained 1000 neurons (to perfom the classification task for 1000 object categories), was replaced with two neurons to adapt the network for performing a binary classification task (i.e. object versus background). The weights of this last fully connected layer were initialized randomly. The network was first trained (all layers, no weight freezing) to perform the CSF versus tissue segmentation task. We then retrained this network to perform microbleed versus tissue segmentation. Training was performed on a single NVIDIA TITAN X with 12 GB GPU memory.

**Figure 3.**
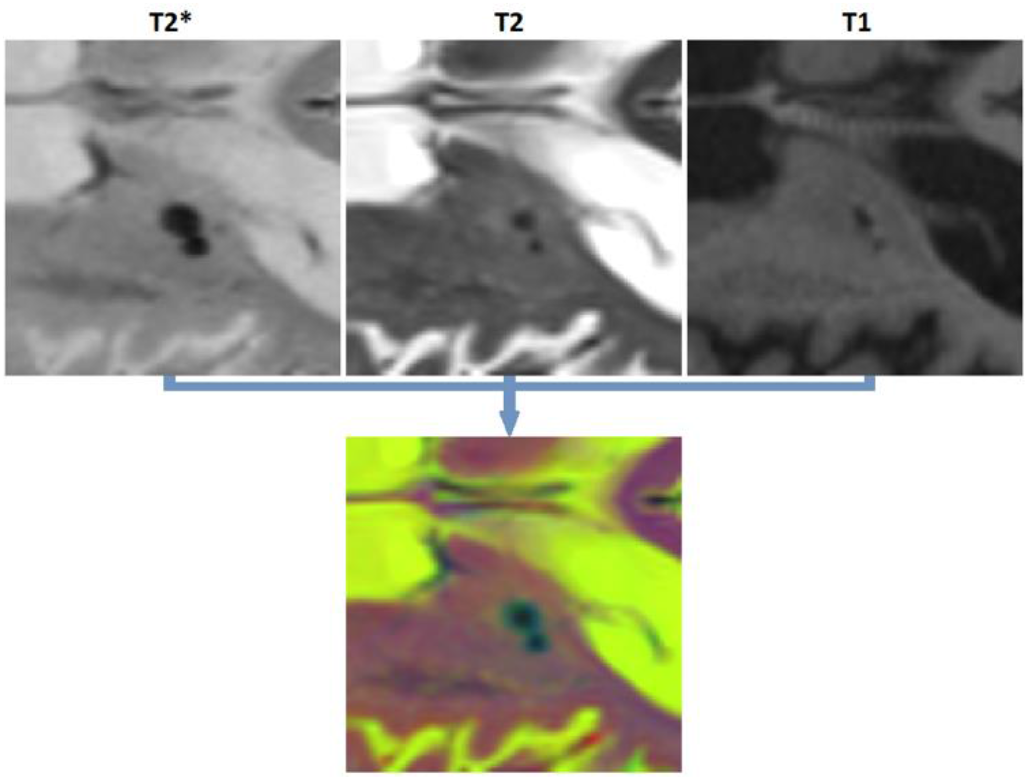
Generating an RGB patch for training ResNet50.

### Parameter Optimization

We assessed the performance of the network with five different patch sizes (14, 28, 52, 56, and 70 mm^2^), with and without freezing the weights of the initial layers (no freezing as well as freezing the first 5, 10, 15, and 20 layers), different mini-batch sizes (20, 40, 60, 80, and 100), and different learning rates (0.002, 0.004, 0.006, 0.008, and 0.010). Each experiment was repeated five times and the results were averaged to ensure their robustness. Epoch parameter was set to 10. Stochastic gradient descent with momentum (SGDM) of 0.9 was used to optimize the cross-entropy loss.

### Post-processing

After applying ResNet50 to segment all microbleed candidate voxels, we performed a post-processing step to reduce the number of false positives as well as to categorize the microbleeds into five classes depending on location. In this post-processing step, the microbleeds were first dilated by two voxels. Then, for each voxel at the border of the microbleed, if the ratio between the T2* intensity of the microbleed voxel and the surrounding dilated area 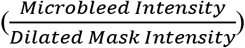 was lower than a specific threshold (to be specified by the user based on sensitivity/specificity preferences), the voxel was removed as a false positive. For each microbleed, the process was repeated iteratively until no voxel was removed as false positive. The final remaining microbleeds were then categorized into regions (cortical GM, deep GM, cerebellum WM, and cerebellum GM and WM based on their overlap with FreeSurfer segmentations). Figure 4 shows examples of FreeSurfer based tissue categories, some segmented microbleeds and dilated surrounding areas.

**Figure 4.**
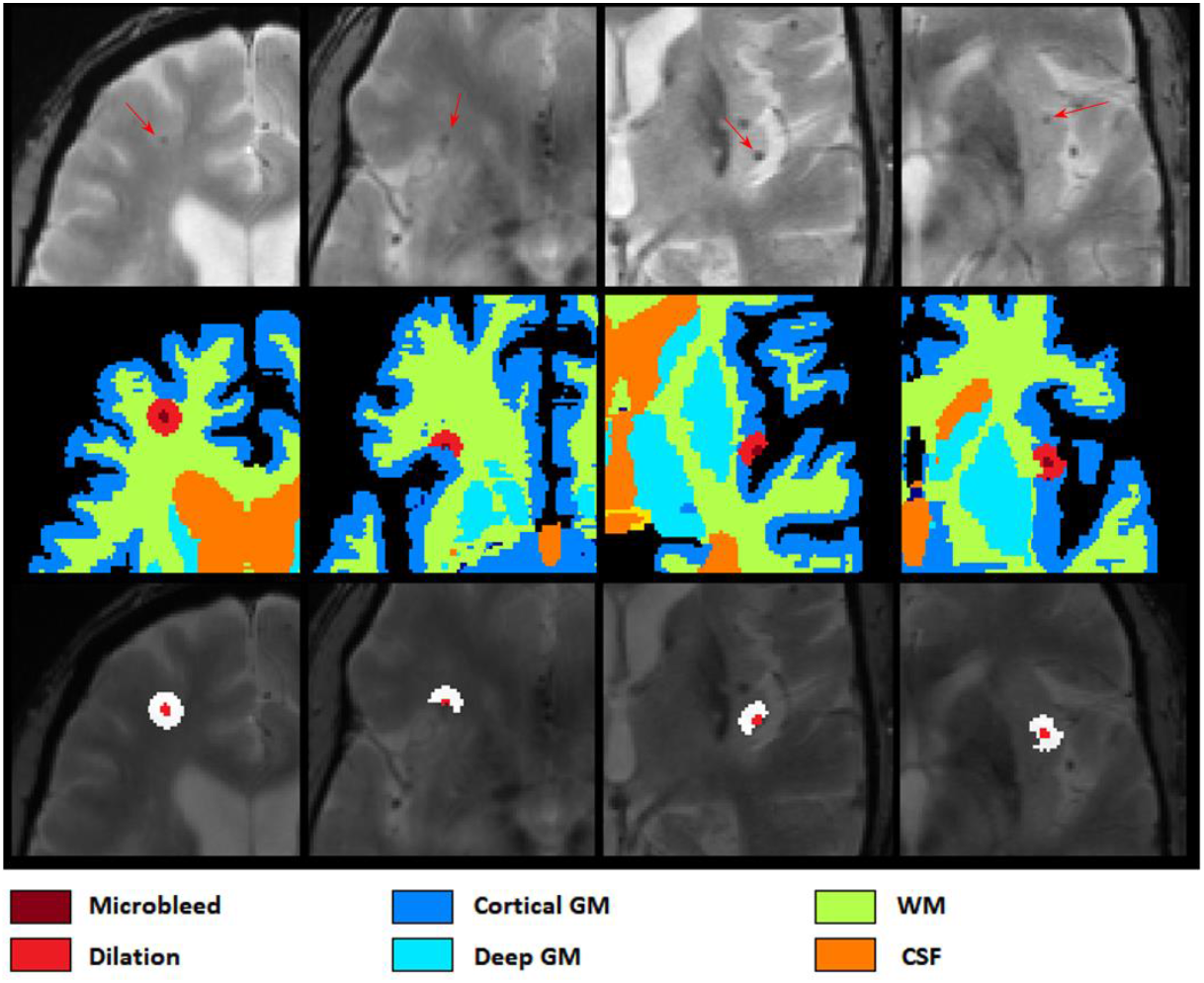
Axial slices showing FreeSurfer based tissue categories, segmented microbleeds and their dilated surrounding areas.

Figure 5 shows a flow diagram of all the different steps performed in the microbleed segmentation pipeline. All implementations (i.e. generating training patches, training and validation of the model, and postprocessing) were performed using MATLAB version 2020b.

**Figure 5.**
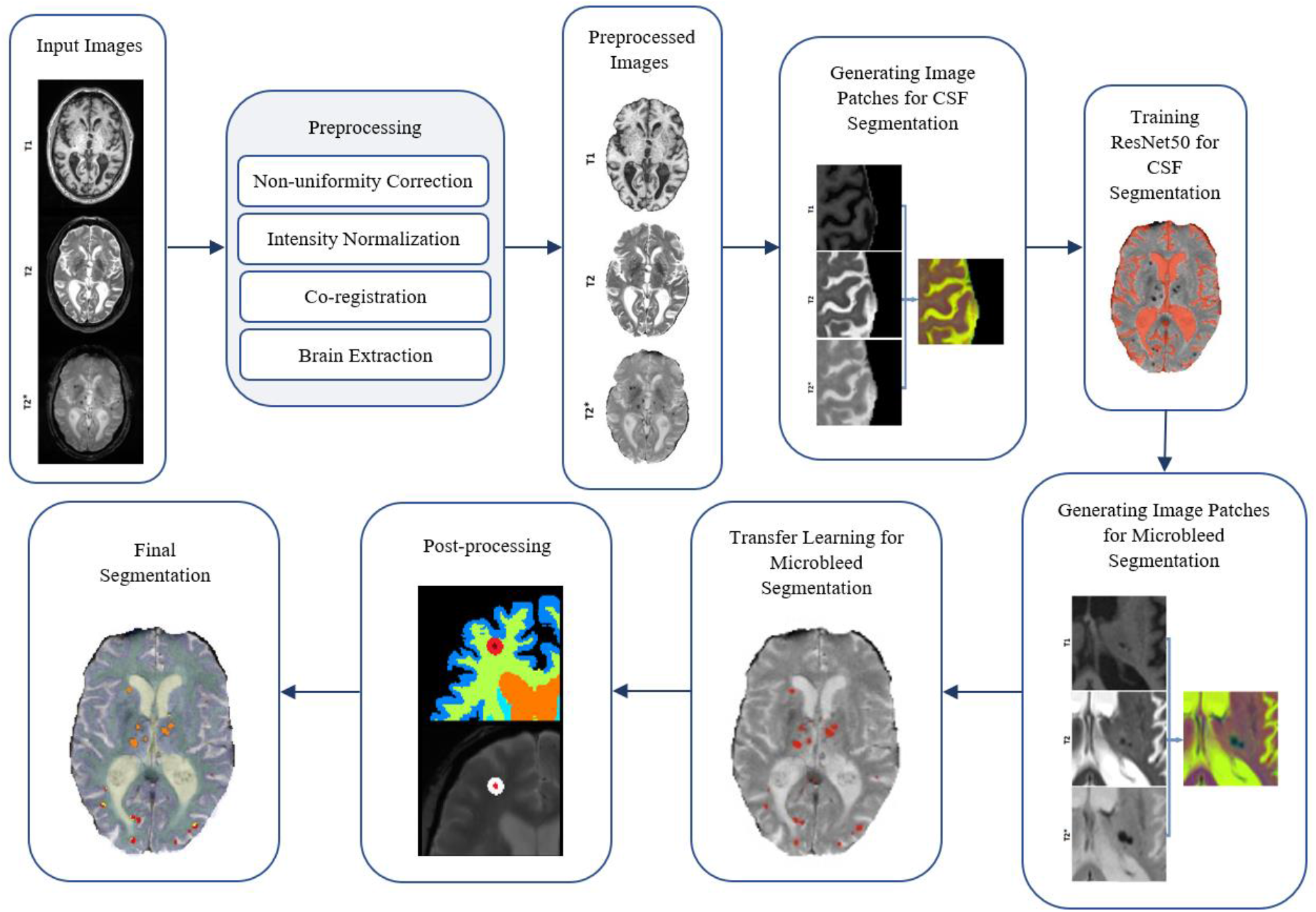
Flow diagram of the microbleed segmentation pipeline.

### Performance Validation

At patch level, to enable comparisons against results of other studies, we measured accuracy, sensitivity, and specificity to assess the performance of the proposed method.

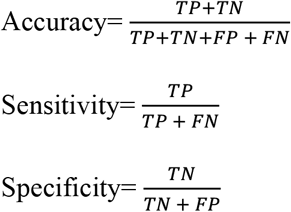

Where TP, TN, FP, and FN denote the number of true positive, true negative, false positive, and false negative patch-level classifications, respectively. While high accuracy at patch-level is desirable, it does not necessarily ensure accurate segmentations at a voxel level. Applied across the entire brain (i.e. 100,000s of pixels), a patch-level specificity of 0.96-0.99% (e.g. as reported by Hong et al. (Hong et al., 2020, 2019)) might translate into thousands of false positive voxels. To assess the performance at a voxel-wise level, we applied the network to all patches in the brain mask, reconstructed the lesions in 3D across patches, and then measured per lesion sensitivity, precision, and Dice similarity index (Dice, 1945) values between manual (considered as the standard of reference) and automated segmentations.

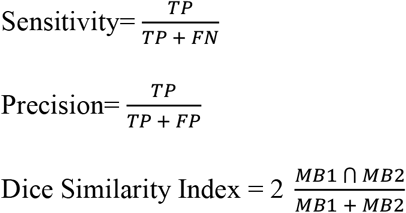

Here, TP (true positive) denotes the number of microbleeds detected by both methods. FN (false negative) denotes the number of microbleeds detected by the manual expert, but not the automated method. FP (false positive) denotes the number of microbleeds detected by the automated method, and not the manual rater. Dice similarity index shows the proportion of microbleeds detection by both methods, over the number of microbleeds detected by each method. A Dice Similarity index of 1 shows perfect agreement between the two methods. A microbleed is considered to be detected by both methods if there are any overlapping microbleed voxels between the two segmentations. Note that in accordance with previous studies, specificity was used to reflect the proportion of true negative versus all negative classifications for patch level results. However, since specificity cannot be defined at per-lesion level, we assessed precision instead of specificity for per-lesion results.

### Data and Code Availability Statement

All image processing steps were performed using minc tools, publicly available at: https://github.com/vfonov/nist_mni_pipelines. BISON (used for tissue classification) is also publicly available at http://nist.mni.mcgill.ca/?p=2148. For more information on the CCNA dataset, please visit https://ccna-ccnv.ca/. The microbleed segmentation methodology has been reported by the inventors to Université Laval (Report on invention 02351) and is now subject to patent protection (U.S. Trademark and Patent Office 63/257, 536).

## RESULTS

### Manual Segmentation and Quality Control

The distribution of manually segmentated lesions for all 78 participants is shown in Figure 6. Based on the manual segmentations, 46.1%, 14.10%, and 58.9% of the cases had at least one microbleed in the cortical GM, deep GM, and WM regions, respectively. Only five cases (6.4%) had microbleeds in the cerebellum (in either GM and WM). The overall intra-rater reliability (per lesion similarity index) for manual segmentation was 0.82 ± 0.14 (κCortical GM = 0.78 ± 0.22, κDeep GM = 1.0 ± 0.0, κWM = 0.81 ± 0.15). All MRIs passed the visual quality control step for preprocessing and BISON/FreeSurfer segmentation.

**Figure 6.**
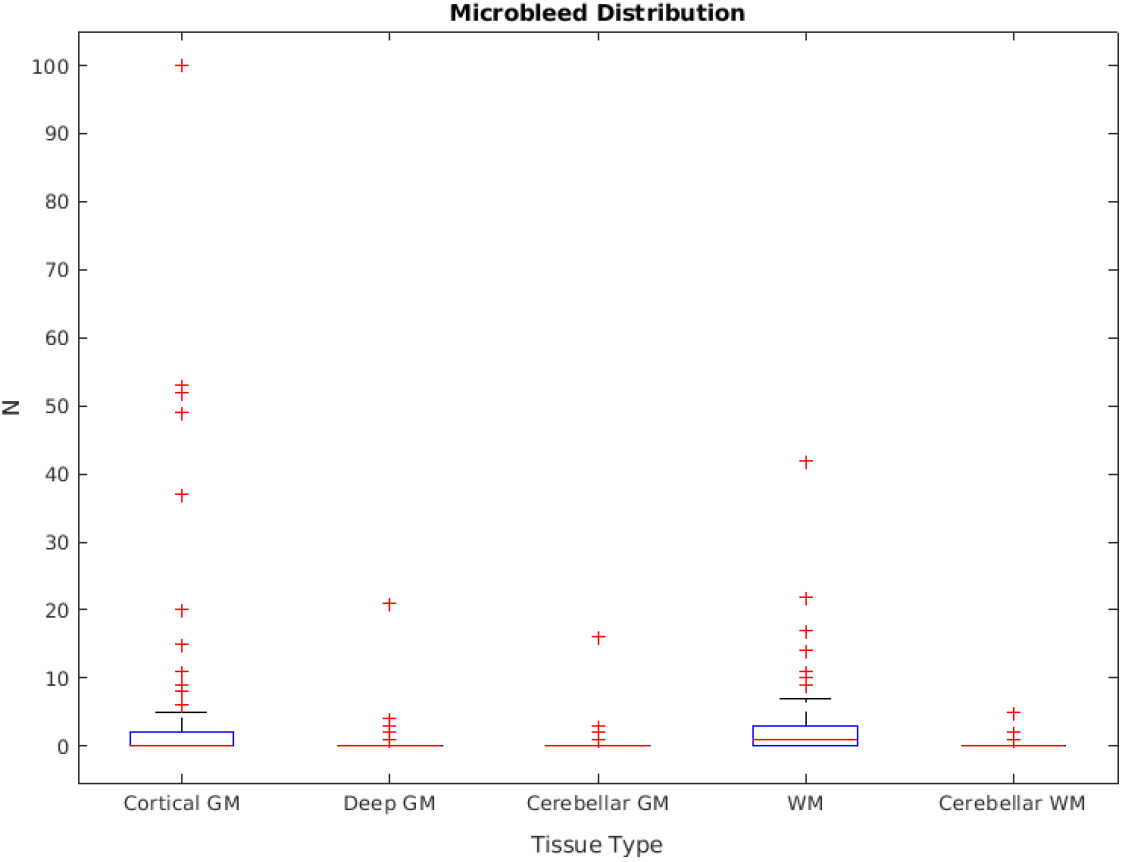
Microbleed distribution for each tissue type, based on the manual segmentations.

### CSF Segmentation

At patch level, ResNet50 segmentations had accuracies of 0.946 (sensitivity = 0.955, specificity = 0.936) and 0.933 (sensitivity = 0.938, specificity = 0.928) for validation and test sets, respecively. At a whole brain voxel level, the segmentations had a Dice similarity index of 0.913 ± 0.015 with BISON segmentations. Overall, ResNet50 CSF segmentations (mean volume = 129.06 ± 31.43 cm^3^) were more generous in comparison with BISON (mean volume = 117.78 ± 29.51 cm^3^). Figure 7 compares the two segmentations, with color yellow indicating voxels that were segmented as CSF with both methods, and colors purple and green indicating voxels that were only segmented as CSF by BISON or ResNet50, respectively. The majority of the disagreements are in the borders of CSF and brain tissue, where ResNet50 segments CSF slightly more generously than BISON.

**Figure 7.**
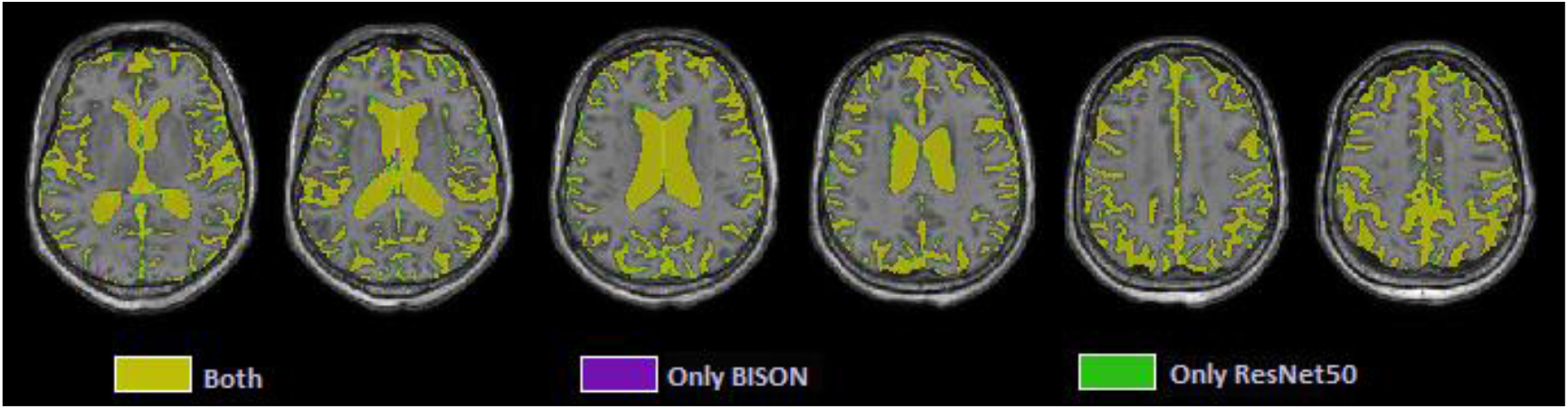
Axial slices comparing ResNet50 and BISON CSF segmentations.

### Microbleed Segmentation

Figure 8 shows the patch level performance results averaged over five repetitions for different patch sizes, freezing of initital layers, mini-batch sizes, and learning rates. Increasing patch size to more than 28 voxel lead to consistently lower accuracy for both validation and test sets. Similarly, learning rates higher than 0.004 lowered the performance. The best performance (in terms of accuracy) was obtained with a patch size of 28, freezing the first five initital layers, a mini-batch size of 40, and a learning rate of 0.006.

**Figure 8.**
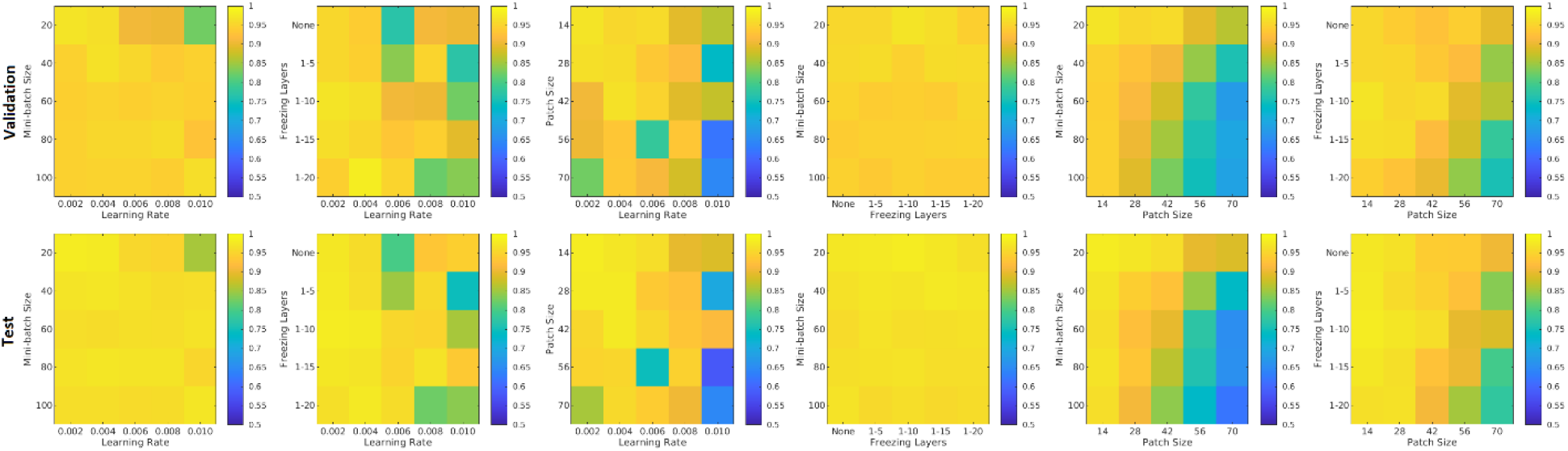
Performance accuracy as a function of patch size, freezing of initital layers, mini-batch size, and learning rate. Colors indicate patch-level accuracy values, with warmer colors reflecting higher accuracy.

Figure 9 shows the average performance of the model with these parameters trained with no augmentation, as well as the same model trained on original data plus data augmented with 4, 9, 14, 19, 24, and 29 random rotations (no augmentation was performed on validation and test sets). All models with data augmentation performed better than the model without any data augmentation. The best performance was obtained using data augmented with 19 random rotations. For this network, patch-level accuracy, sensitivity, and specificity values were respectively 0.990, 0.979, and 0.999 for the validation set, and 0.996, 0.992, and 0.999 for the test set.

**Figure 9.**
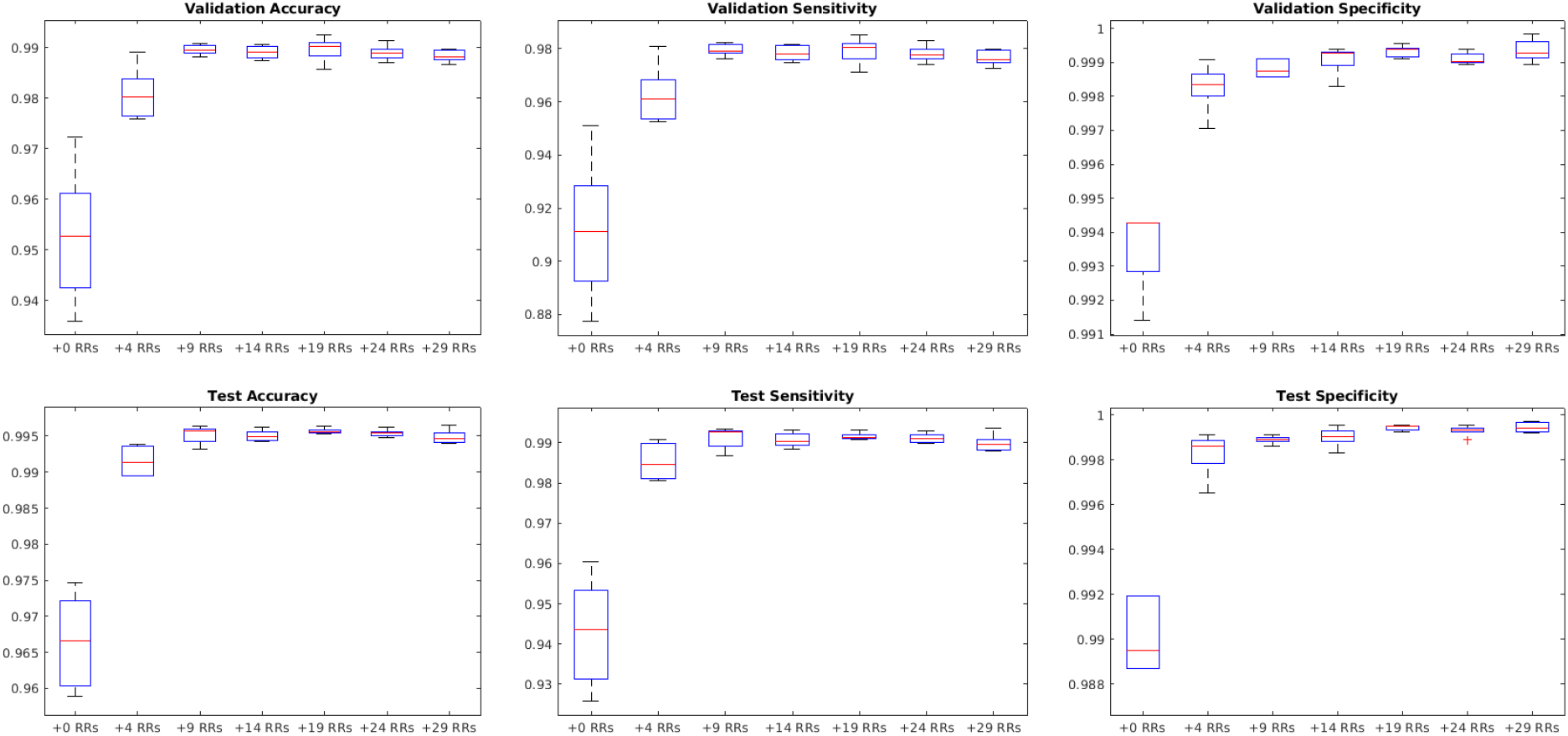
Impact of data augmentation on performance accuracy, sensitivity and specificity at patch level. RR= Random Rotation.

Figure 10 shows sensitivity, precision, and similarity index values separately for cortical, deep, and cerebellar GM, and cerebral and cerebellar WM, after applying post-processing with different thresholds to the voxel-wise segmentations. The threshold values can be selected by the user based on sensitivity and precision preferences.

**Figure 10.**
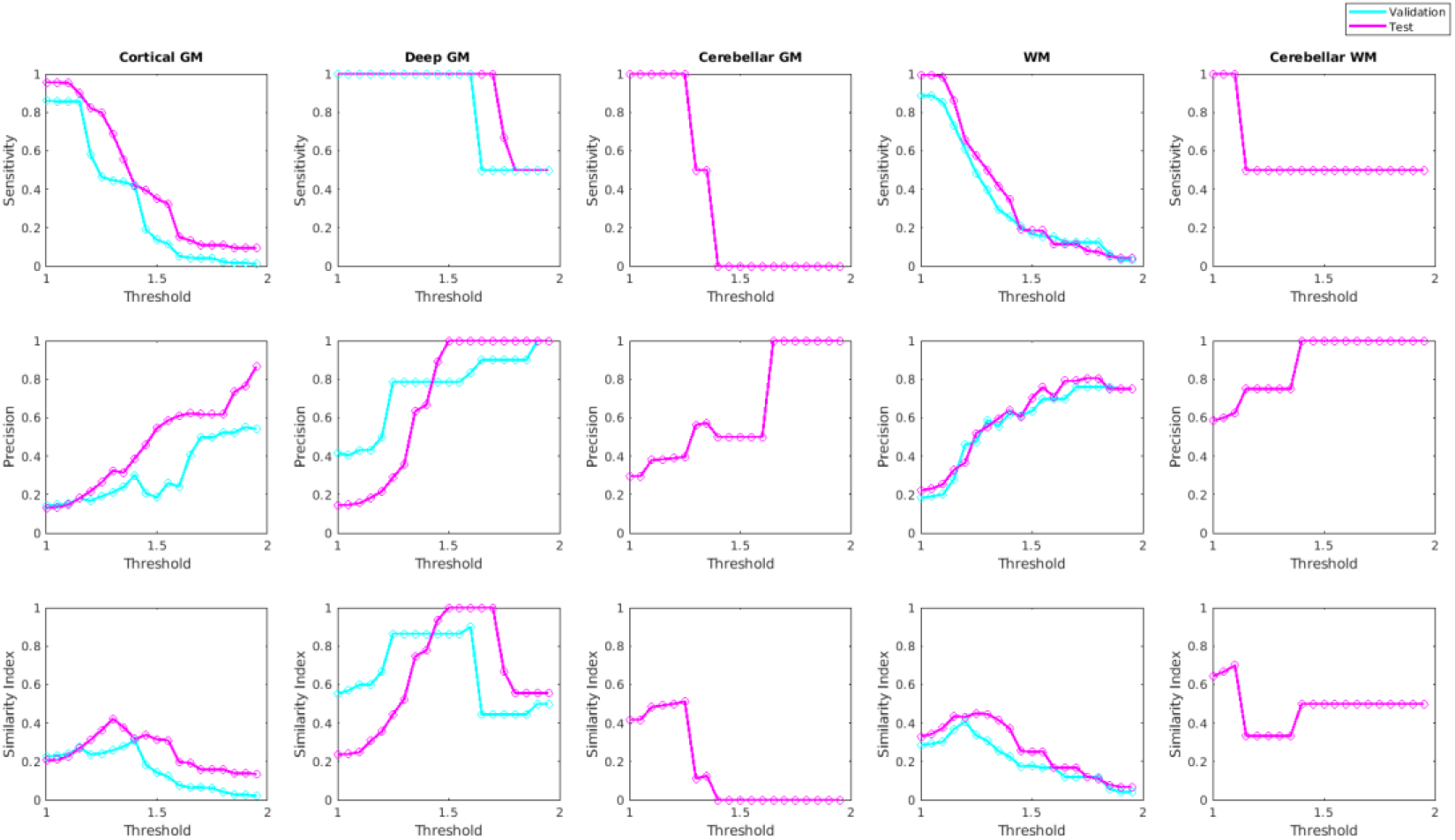
Sensitivity, precision, and similarity index values for different post-processing threshold values in validation and test sets.

Figure 11 shows examples of automated versus manual segmentations for threshold = 1.4 for the deep GM and 1.2 for the rest of the regions, with examples of true positive (indicated in red), false positive (indicated in blue), and false negative (indicated in green) classifications. Most of the disagreements are in the voxels in the border of the microbleeds, where the automated tool sometimes performs a more generous (i.e. blue voxels) or more conservative (i.e. green voxels) segmentation than the manual expert. Such differences will not affect the overal microbleed counts, since both methods have essentially identified the same microbleed.

**Figure 11.**
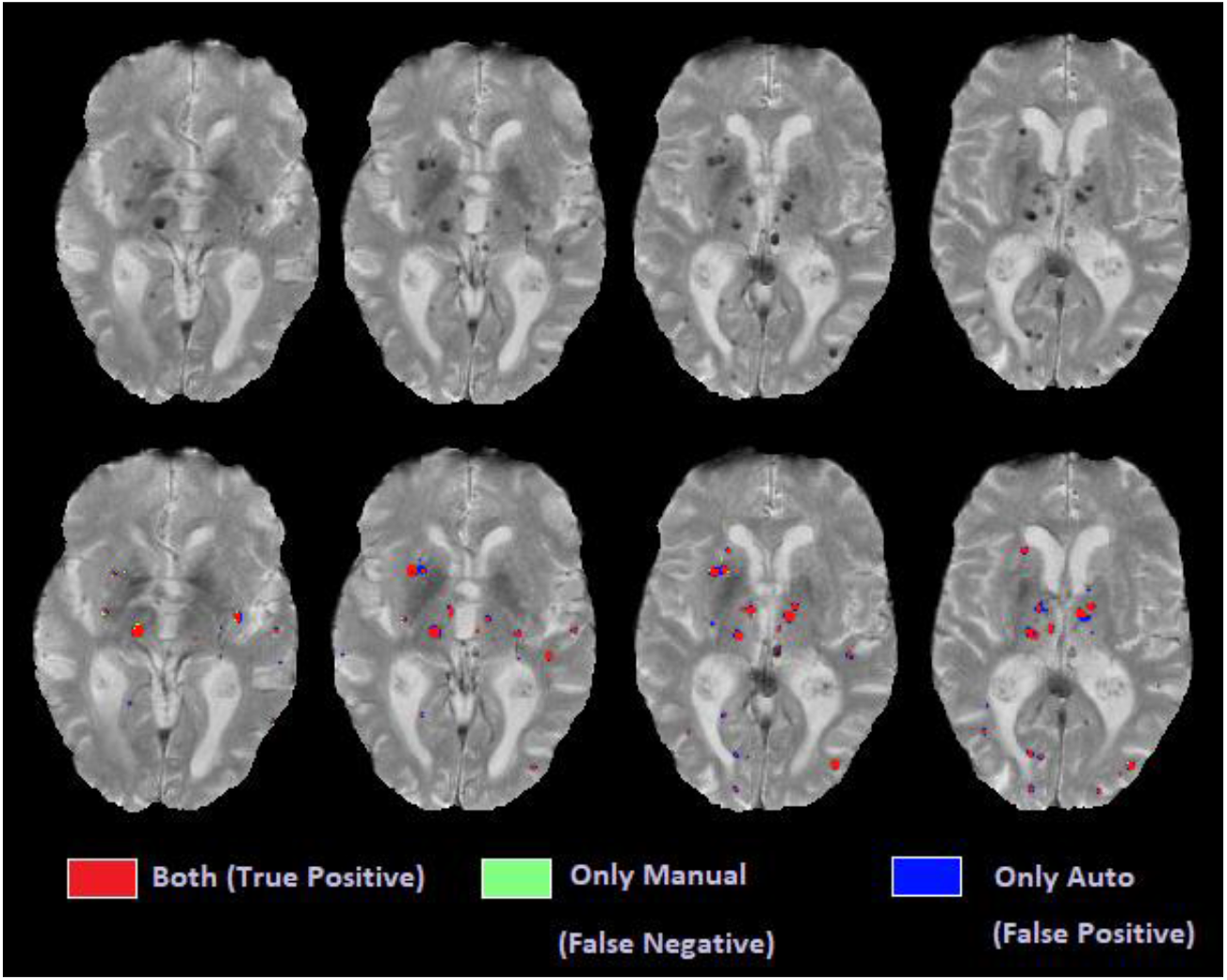
Axial slices comparing automated and manual segmentations.

## DISCUSSION

In this paper, we presented a multi-sequence microbleed segmentation tool based on ResNet50 network and routine T1-weighted, T2-weighted, and T2* MRI. To overcome the lack of availability of a large training dataset, we used transfer learning to first train ResNet50 for the task of CSF segmentation, and then retrained the resulting network to perform microbleed segmentation.

The CSF classification experiments showed excellent agreement (Dice similarity index = 0.91) between ResNet50 and BISON segmentations. The majority of the disagreements were in the voxels bordering the CSF and brain tissue, where ResNet50 segmented CSF slightly more generously than BISON. Given that BISON segmentations were based only on T1-weighted images, whereas ResNet50 used information from T1-weighted, T2-weighted, and T2* sequences, these voxels might actually be CSF voxels correctly classified by ResNet50 method that did not have enough contrast on T1-weighted images to be captured by BISON.

At patch level, with sensitivity and specificity values of 99.57% and 99.93%, the microbleed segmentation method outperforms most previously published results. At a per lesion level, the strategy yielded sensitivity, precision, and Dice similarity index values of 89.1%, 20.1%, and 0.28 for cortical GM, 100%, 100%, and 1 for deep GM, and 91.1%, 44.3%, and 0.58 for WM, respecitvely. Post-processing improved the results (increased the similarity index) for all microbleed types (by successfully removing the false positives). The improvement was most evident for deep GM microbleeds, where most of the false positives were at the boundaries of the deep GM structures, which tend to have lower intensities compared to the neighboring tissue.

There are inherent challenges in comparing the performance of our proposed method against previously published results. Other work has been mostly based on susceptibility-weighted scans, which in general have higher sensitivity and resolution levels (usually acquired at 0.5×0.5 mm^2^ voxels versus 1×1 mm^2^ voxels in our case) and are better suited for microbleed detection (Dou et al., 2016; Hong et al., 2020; Roy et al., 2015; Shams et al., 2015; Van Den Heuvel et al., 2016; Wang et al., 2017; Zhang et al., 2018). Another concern in comparing results across studies regards the characteristics of the dataset used for training and validation of the results. Most previous methods have used data from populations that are much more likely to have microbleeds, such as patients with cerebral autosomal-dominant arteriopathy with subcortical infarcts and leukoencephalopathy (also known as CADASIL) (Hong et al., 2020; Wang et al., 2017; Zhang et al., 2018). In contrast, our dataset included non-demented aging adults and patients with neurodegenerative dementias which do not necessarily have such a high cerebrovascular disease burden. In fact, 63% of the cases in our sample had fewer than three microbleeds. Since we use a participant-level assignment in training, validation, and test sets, even one false positive would reduce the per-participant precision value by 33-50% for those cases. In comparison, the training dataset used by Dou et al. included 1149 microbleeds in 20 cases (i.e. 57.45 microbleeds per case on average), in which case one false positive detection would only change the reported precision by 1.7% (Dou et al., 2016). Along the same line, we included very small microbleeds with volumes between 1-4 mm^3^ (i.e. 1 to 4 voxels) in our sample (∼35% of the total microbleed count, distributed consistently between training, validation, and test sets), whereas others might choose to not include such very small lesions which are more challenging to detect and also have lower inter and intra rater reliability (Cordonnier et al., 2007). Regardless, even considering the disadvantage of fewer microbleeds per scan inherent to our population, the proposed method compares favorably against other published results.

Generalizability to data from other scanner models and manufacturers is another important point when developing automated segmentation tools. Automated techniques that have been developed based on data from a single scanner might not be able to reliably perform the same segmentation task when applied to data from different scanner models and with different acquisition protocols (Dadar and Duchesne, 2020). To ensure the generalizability of our results, we used data from seven different scanner models across three widely used scanner manufacturers (i.e. Siemens, Philips, and GE) from a number of different sites.

Due to the inherently difficult nature of the task, even in manual ratings, inter-rater and intra-rater variability in microbleed detection is generally not very high. The per-lesion intra-rater similarity index for our dataset (based on eight randomly selected cases) was 0.82. Other studies have also reported similar results, with one study reporting intra-rater and inter-rater agreements (similarity index) of 0.93 and 0.88 respectively, while others generally report more modest inter-rater agreements ranging between 0.33-0.66 (Cordonnier et al., 2007). In a dataset of 301 patients and using T2* images for microbleed detection, Gregoire et al. reported inter rater similarity index values of 0.68 for presence of microbleeds, where the two raters detected 375 (range: 0–35) and 507 (range: 0–49) microbleeds respectively (Gregoire et al., 2009). Given the relatively high levels of inter-rater and intra-rater variability in microbleed segmentation results, it is also possible that some of the false positives detected by the automated method might be actual microbleed cases that were missed by the manual rater. Regarding the location of the microbleeds, Gregoire et al. reported higher levels of agreement (between the manual ratings) for microbleeds in the deep GM regions, similar to our results. This could be explained by the higher intensity contrast between the microbleeds (greater levels of hypointensity) and the background GM, which has a higher T2* signal than the WM, where the performance is usually less robust for manual raters as well (Gregoire et al., 2009). Finally, the relatively lower performance for cortical GM microbleed segmentation is also expected, given the close proximity to blood vessels (both in the sulci and on the surface of the brain), which show a hypointense signal that confounds with that of the microbleeds, leading to false positiveas and lowering the precision. However, despite the different levels of performance in the different brain areas, an automated segmentation method has the clear advantage of providing robust segmentations across different runs, essentially eliminating intra-rater variability.

Accurate and robust microbleed segmentation is necessary for assessing cerebrovascular disease burden in the aging and neurodegenerative disease populations, who may show a lower number of microbleeds than other pathologies (e.g. CADASIL), making the task more challenging. Additionally, an automated tool that can reliably detect microbleeds using data from different scanner models is highly advantageous. Our results suggest that the proposed method can provide accurate microbleed segmentations in multi-scanner data of a population with a low number of microbleed per scan, making it applicable for use in large multi-center studies.

## ACKNOWLEDGEMENTS

MD is supported a scholarship from the Canadian Consortium on Neurodegeneration in Aging in which SD and RC are co-investigators as well as an Alzheimer Society Research Program (ASRP) postdoctoral award. The Consortium is supported by a grant from the Canadian Institutes of Health Research with funding from several partners including the Alzheimer Society of Canada, Sanofi, and Women’s Brain Health Initiative.

